# HER4 is a high affinity dimerization partner for all EGFR/HER/ErbB-family proteins

**DOI:** 10.1101/2024.05.03.592409

**Authors:** Pradeep Kumar Singh, Soyeon Kim, Adam W. Smith

## Abstract

Human epidermal growth factor receptors (HER) – also known as EGFR or ErbB receptors – are a subfamily of receptor tyrosine kinases (RTKs) that play crucial roles in cell growth, division, and differentiation. HER4 (ErbB4) is the least studied member of this family, partly because its expression is lower in later stages of development. Recent work has suggested that HER4 can play a role in metastasis through cell migration and invasiveness; however, unlike EGFR and HER2, the precise role that HER4 plays in tumorigenesis is still unresolved. Early work on HER family proteins suggested that there are direct interactions between the four members, but to date, there has been no single study of all four receptors in the same cell line studied with the same biophysical method. Here, we quantitatively measure the degree of association between HER4 and the other HER-family proteins in live cells with a time-resolved fluorescence technique called pulsed interleaved excitation fluorescence cross-correlation spectroscopy (PIE-FCCS). PIE-FCCS is sensitive to the oligomerization state of membrane proteins in live cells, while simultaneously measuring protein expression levels and diffusion coefficients. Our PIE-FCCS results demonstrate that HER4 interacts directly with all HER family members in the cell plasma membrane. The interaction between HER4 and other HER family members intensified in the presence of a HER4-specific ligand. Our work suggests that HER4 is a preferred dimerization partner for all HER family proteins, even in the absence of ligands.

## Introduction

Human epidermal growth factor receptor (HER)-family proteins play critical roles in development and homeostasis, but they can also drive severe health issues when mutated or overexpressed(Appert-Collin et al. 2015; Tebbutt et al. 2013). There are four members of the HER family, including EGFR (HER1/ErbB1), HER2 (ErbB2), HER3 (ErbB3), and HER4 (ErbB4). EGFR and HER4 are classic receptors with a ligand-binding extracellular domain and an intracellular catalytic domain, while HER2 has no known ligand and HER3 is kinase-deficient(Citri et al. 2003; Lemmon et al. 2014; Riese et al. 1996; Walker 1998). Generally, ligand binding to HER proteins leads them to dimerize with themselves or other related family members, and dimerization promotes tyrosine phosphorylation(Citri and Yarden 2006; Endres et al. 2014; Yarden and Sliwkowski 2001). The kinase activity of HER4 is linked to MAPK and (PI3K)/AKT pathways as well as other downstream signaling events(Carpenter 2003; El-Gamal et al. 2021). A wide range of adapters and signaling proteins dock with these phosphorylated tyrosine residues(Carraway and Sweeney 2001; Hynes and MacDonald 2009). Unregulated tyrosine kinase activity associated with HER family members may promote tumorigenesis in breast, lung, and colon cancer and several monoclonal antibodies and tyrosine kinase inhibitors (TKIs) that target HERs have been approved by the FDA to treat cancer patients(Fujiwara et al. 2014; Hynes and MacDonald 2009; Kumagai et al. 2021; Tebbutt et al. 2013).

Among HER-family receptors, EGFR and HER2 are arguably the most well-studied because of their strong association with various cancers of the lung, brain, and breast. HER3 is a frequent dimerization partner with HER2, and dual anti-HER2/anti-HER3 therapy has demonstrated efficacy in the treatment of breast cancer(Blumenthal et al. 2013; Diwanji et al. 2021; Phillips et al. 2014). In contrast, HER4 is not as commonly associated with cancer. The HER4 protein was first discovered by Plowman et al. in MDA-MB-453 cells while searching for specific ligands for other membranes of the HER family(Plowman et al. 1993). HER4 protein expression is high in fetal cardiac muscle, brain, and testis, while low in adult tissues, which supports its role in differentiation and development. More recent work has reported its presence in all adult tissues apart from glomeruli and peripheral nerves(Muraoka-Cook et al. 2008; Qiu et al. 2008), however the expression levels are generally lower than the other HER family members. There are three structural regions in HER family proteins: an extracellular domain (ECD), a transmembrane domain (TMD), and an intracellular domain (ICD)(Sweeney et al. 2000). The HER4 ECD is most similar to HER3, while its cytoplasmic domain exhibits 79% homology with EGFR and 77% with HER2. Despite 60-79% sequence identity, HER proteins differ in many ways beyond their specificity for ligands and effectors. EGFR and HER2 lack phosphorylation sites that allow PI3K subunit p85 interaction; however, HER3 and HER4 contain phosphorylation sites that permit direct PI3K signaling(Yarden and Sliwkowski 2001). Similarly, EGFR phosphorylation can bind directly to ubiquitin ligase CBL, while HER4 requires the adaptor protein GRB2 for CBL binding(Lucas et al. 2022). As a result of these structural differences, EGFR and HER2 tend to increase cell proliferation, while HER4 generally suppresses cell proliferation(Muraoka-Cook et al. 2008; Xu et al. 2018). Despite this canonical view, many reports have highlighted the potential role of HER4 in cancer development, but the context dependence of HER4’s oncogenic role is still not fully understood(Lucas et al. 2022; Roskoski 2014).

Heterodimerization is an essential step for the activation of catalytically impaired receptor HER3 and the orphan receptor HER2, and many experimental studies of have reported on direct heterodimerization between HER2 and HER3(Citri et al. 2003; Roskoski 2014; Tao and Maruyama 2008; Weitsman et al.). More recently, structural studies have resolved the interactions between HER2 and EGFR(Bai et al. 2023; Diwanji et al. 2021). Functional studies have explored the effects of EGFR or HER2 on HER4 activation(Yarden and Sliwkowski 2001). For example, kinase-dead HER4 mutants were found to be as efficient as wild-type HER4 in forming a heterodimeric assembly with HER2(Graus-Porta et al. 1997). Many biophysical techniques have been developed and applied to resolving heterodimers between HER proteins to investigate their role in biology(Brown et al. 2022; Garrett et al. 2003; Nagy et al. 2010; Ogiso et al. 2002; Pryor et al. 2015; Steinkamp et al. 2014; Tao and Maruyama 2008). High resolution structure approaches provide atomic level details, but quaternary interactions between membrane proteins need to be resolved in situ because of the chemical complexity of the plasma membrane. Several fluorescence-based methods have been developed to quantify protein interactions in live cells(Martin-Fernandez 2023; Sankaran and Wohland 2023; Stoneman and Raicu 2023). Fluorescence fluctuation spectroscopy (FFS) methods have become valuable for analyzing membrane protein interactions(Bacia et al. 2006; Christie et al. 2020). FFS offers insight into temporal and spatial dimensions that are not easily accessible by super-resolution approaches(Christie et al. 2020). One of these FFS methods is pulsed interleaved excitation fluorescence cross-correlation spectroscopy (PIE-FCCS), which is specialized for a multi-parameter characterization of membrane protein interactions in living cells(Christie et al. 2020; Müller et al. 2005).

Our lab has used PIE-FCCS to investigate conformational coupling across the cell membrane and the multimeric structure of EGFR activated by ligands(Endres et al. 2013; Huang et al. 2016). In addition, we have been able to resolve the clinical implications of oncogenic mutants of EGFR using this approach(Brown et al. 2022; Du et al. 2021). The PIE-FCCS technique can be used to determine membrane protein expression levels, diffusion coefficients, and the degree of cross-correlation (abbreviated as *f*_*c*_), which is a direct measure of how strongly the proteins interact and diffuse together(Christie et al. 2020). It does not detect immobile aggregates or internalized proteins, as it is primarily sensitive to diffusing proteins. As the degree of association depends on correlated diffusion, we can interpret the co-diffusing species as stable over the timescale of the transit time (∼10^−1^ s), although it is not possible to directly resolve association lifetimes.

In this work, we use PIE-FCCS to evaluate multimerization between HER-family proteins and examine the effects of different ligand stimulation conditions. The measurements provide direct vidence for membrane protein interactions in live cells at physiological expression levels. Overall, we find that HER4 does not self-associate prior to ligand binding, however, it does interact with all other HER family members. The degree of HER4 heterodimerization increases upon NRG1 stimulation but is unaffected by EGF treatment. Heterodimerization between HER4 and all other HER-family proteins also increases upon NRG1 stimulation. Our findings suggest that HER4 is a high-affinity heterodimer partner for all of the other HER-family proteins.

## Results

### In resting cells, EGFR, HER2, and HER4 are homo-monomers, but HER3 is a homodimer

We first set out to measure the degree of multimerization for each HER-family protein. We expressed each protein as two C-terminal fluorescent protein fusion constructs (i.e. HER-mCherry and HER-eGFP) in COS-7 cells and quantified the interactions using PIE-FCCS (Fig. 1A). Representative epifluorescence images are shown for COS-7 cells expressing mCherry and eGFP tagged EGFR, HER2, HER3 and HER4 (Fig. S1). The optical layout for the PIE-FCCS instrument is outlined in Fig. 1A along with a representative COS-7 cell with HER4 protein expression and a schematic of membrane protein diffusion. Fig. 1B displays a representative set of single-cell PIE-FCCS data for HER4 proteins. The red and green autocorrelation functions (ACF) are fit to determine the density and mobility of mCherry- and eGFP-tagged HER4 respectively. The blue line shows the cross-correlation function (CCF), the amplitude of which is indicative of correlated diffusion. The fitted values, *f*_*c*_ and D_eff_, for each single cell measurement are summarized in Fig. 1C as boxplots. The distribution of *f*_*c*_ values for EGFR had a median value of 0.01 (Fig. 1C, green). EGFR showed an average diffusion coefficient of 0.31 μm^2^/s in the resting cell environment (Fig. 1C, green). The expression levels of the EGFR protein were calculated to be between 100 to 2000 receptors/μm^2^.

**Figure 1.**
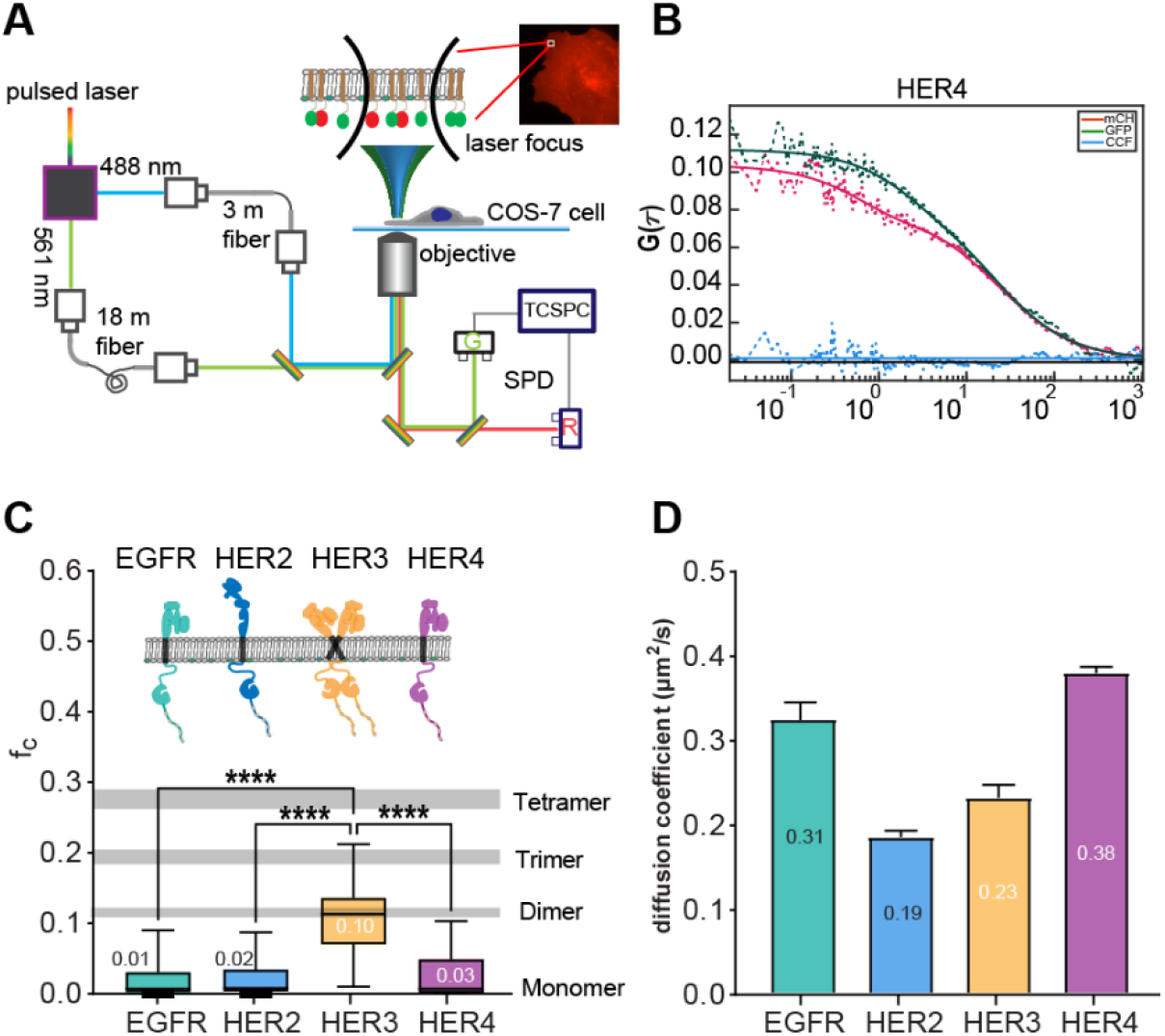
(A) Schematic of the PIE-FCCS instrument with two-color laser excitation. The inset shows an epifluorescence image of a COS-7 cell expressing HER4-eGFP. A full description of the PIE-FCCS instrument is given in the method section. (B) A representative set of single-cell PIE-FCCS data is shown, with the two ACF in green and magenta and the CCF in blue. (C) The distribution of *f*_*c*_ values is shown for all of the single cell homodimerization measurements of each HER protein without ligand addition. The distributions are represented as boxplots with the median value listed beside each box. The boxes enclose the 25-75 percentile range and the whiskers show the entire range of data. (D) Diffusion coefficients are extracted from the ACF data for each of the measurements shown in panel C. The data is plotted as the mean of the distribution ± the standard error (SEM).

Several controls have been used to interpret the PIE-FCCS results. First, a solution of duplex DNA (Fig. S2, T4-DNA, red) was used for laser alignment. Second, membrane protein constructs Src and GCN4 were used as live-cell monomer and dimer controls, respectively (Fig. S2, S3). A detailed and comprehensive explanation of the control samples and the precise process of extracting the *f*_*c*_ value has been given in the method section and in previous work from our group(Kaliszewski et al. 2018).

The PIE-FCCS results for HER2 expressing cells also show near-zero *f*_*c*_ values (median *f*_*c*_ = 0.02, Fig. 1B, blue), indicating that it does not homodimerize significantly in resting cells. Compared with EGFR, HER2 receptors exhibit a lower diffusion coefficient (0.19 μm^2^/s, Fig. 1C, blue). The cross-correlation results are consistent with previous reports that HER2 does not homodimerize in resting cells(Diwanji et al. 2021; Graus-Porta et al. 1997). While surprising, the low diffusion coefficient of HER2 is likely due to interactions with other membrane proteins including HER3(Jaulin-Bastard et al. 2001; Jeong et al. 2017). We have measured HER2 and HER3 heterodimerization with PIE-FCCS and summarized the results in Fig. S10A-B. The median *f*_*c*_ value of 0.18 shows that HER2 and HER3 proteins form a strong heterodimer prior to NRG1 ligand binding. This ligand independent heterodimerization of HER2 and HER3 has been reported in several earlier studies(Diwanji et al. 2021; Pryor et al. 2015; Steinkamp et al. 2014; Weitsman et al.).

PIE-FCCS measurements of HER3 revealed a median *f*_*c*_ = 0.10, which indicates significant homodimerization (Fig. 1C, yellow). The average diffusion coefficient of HER3 is 0.23 um^2^/s (Fig. 1D, yellow), which is slower than monomeric EGFR and consistent with ligand independent dimerization. This result is somewhat surprising as early studies reported that HER3 was incapable of homodimerization(Berger et al. 2004). However, our results are consistent with more recent single molecule imaging, which provided evidence of HER3 multimerization(Pryor et al. 2015; Steinkamp et al. 2014).

Like EGFR and HER2, HER4 showed near-zero cross-correlation (median *f*_*c*_ = 0.03, Fig. 1C, magenta), indicating that the proteins are primarily monomeric in COS-7 cells. To the best of our knowledge, no investigation has reported on the dimerization/oligomerization state of HER4 in the absence of ligand stimulation. The diffusion coefficient of HER4 is 0.38 μm^2^/s, (Fig. 1D, magenta) which is the highest in comparison to other HER family members, but similar in magnitude to EGFR. Single cell PIE-FCCS data of all four homomeric interactions (Fig. S4), and fit parameters (Table S1) can be found in the supplemental information document. Overall, these PIE-FCCS measurements demonstrate that EGFR, HER2, and HER4 do not self-associate in the absence of ligand, while HER3 exists as a homodimer.

### HER4 forms a dimer with NRG1 and is unaffected by EGF stimulation

We next tested the effect of ligand stimulation with two common ligands for HER-family proteins: EGF and NRG1(Dawson et al. 2007; Nagy et al. 2010; Plowman et al. 1993). COS-7 cells were transiently transfected with HER4-eGFP and HER4-mCherry and data were collected at cell surface densities between 100-1200 molecules/μm^2^. We determined the fraction of correlation and diffusion coefficients using the PIE-FCCS data. EGF is not a natural ligand for HER4 and so as expected, EGF stimulation (500 ng/ml) does not change the oligomeric state of HER4 (*f*_*c*_ = 0.03), as shown in Fig. 2A (blue). The diffusion coefficient of the protein did not change with EGF ligand stimulation, consistent with the *f*_*c*_ values (Fig. 2B, 2C, grey and blue). Upon the addition of NRG1, there was a significant increase in cross-correlation (median *f*_*c*_ = 0.14), indicating the formation of a homodimer complex (Fig. 2A, magenta; Fig. S2B). The HER4 *f*_*c*_ distribution matches the value obtained from a known membrane protein dimer (e.g. GCN4 Fig. S2B) and the numerical value expected for dimerization in stochastic model(Kaliszewski et al. 2018). Based on these comparisons we will refer to these complexes as HER4 homodimers. The diffusion coefficient of HER4 fell after ligand addition, which supports the interpretation of the *f*_*c*_ values (Figs. 2B and 2C, magenta). Single cell PIE-FCCS data of all four homomeric interactions (Fig. S5), and fit parameters (Table S2) can be found in the supplemental information document.

**Figure 2.**
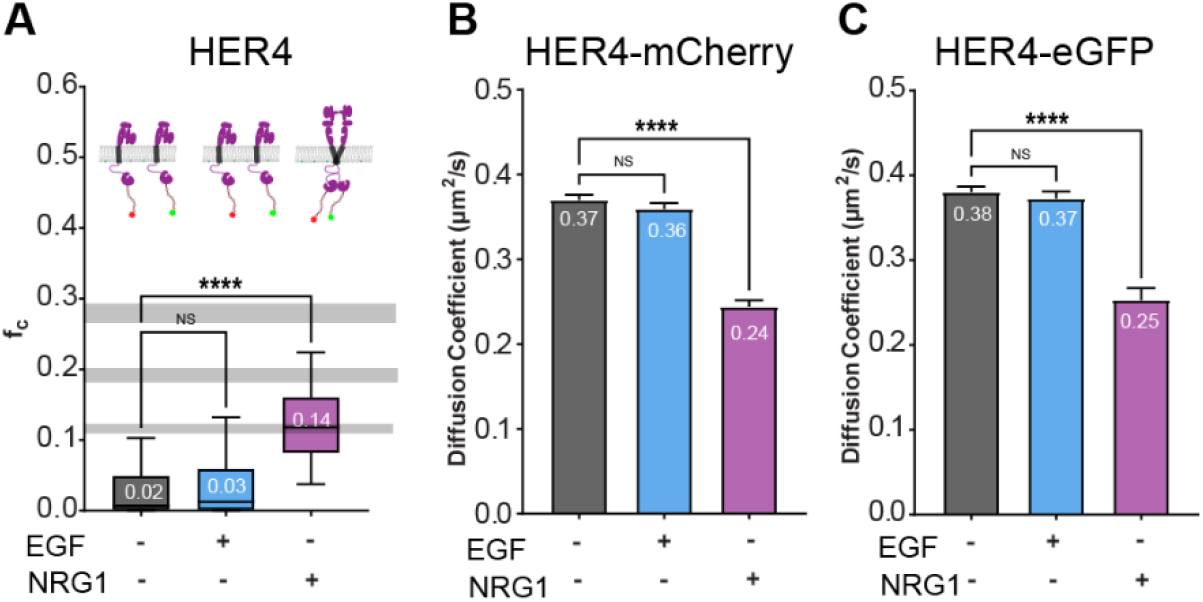
Summary of PIE-FCCS measurements of HER4 before and after ligand stimulation. Panel (A) shows the distribution of single cell *f*_*c*_ values for HER4 homodimerization without ligand stimulation and with EGF or NRG1 addition. An illustration of the HER4 dimerization state is shown above each set of data. The numbers within each box represent the median *f*_*c*_ values. The boxes enclose the 25-75 percentile and the whiskers enclose the entire range of data. Panels (B & C) represent the average diffusion coefficients of each protein before and after ligand treatment. Generally, diffusion coefficients will go down when *f*_*c*_ values go up as larger multimers will diffusion slower than smaller multimers. Diffusion coefficient data is mean values ± SEM.

### HER4-EGFR forms a dimer in resting cells and a multimer with ligand (NRG1) stimulation

To quantify the interactions between HER4 and EGFR, we co-expressed them in live cells and measured the degree of association with PIE-FCCS. Single cell PIE-FCCS data and fit parameters can be found in Fig. S6 and Table S3), and the results are summarized in Fig. 3. Interestingly, HER4 and EGFR have a median *f*_*c*_ value of 0.12 without ligand addition, indicating that they heterodimerize significantly in resting cells (Fig. 3A, grey). The effect of ligand stimulation on heteromeric interactions was investigated using two ligands (EGF and NRG1). COS-7 cells stably co-expressing HER4-mCherry and EGFR–eGFP were stimulated with 500 ng/ml EGF or 500 ng/ml NRG for 15 minutes and subsequently used for PIE-FCCS measurements. In the presence of EGF, the HER4-EGFR cross-correlation increases slightly (*f*_*c*_ = 0.17, Fig. 3A, blue), indicating that the ligand promotes heterodimerization, but only to a small degree. Addition of NRG1 ligand nearly doubles the median *f*_*c*_ values from 0.12 to 0.21 (Fig. 3A) indicating heteromultimerization of HER4 and EGFR.

**Figure 3.**
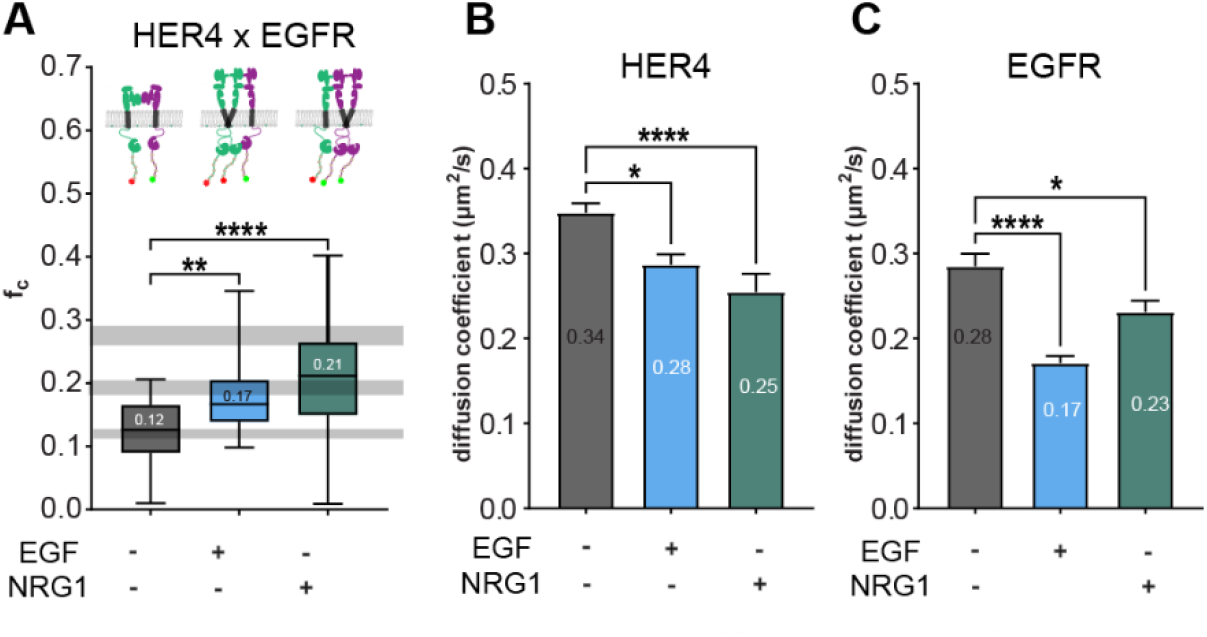
PIE-FCCS measurement of HER4-EGFR heteromer with ligands stimulation. Panel (A) shows the degree of cross-correlation (*f*_*c*_) between HER4 and EGFR proteins before (-) and after (+) EGF and NRG1 stimulation. An illustration of the multimerization states is shown above each set of data. Panels (B & C) represent the average diffusion coefficients of each protein (HER4 and EGFR, respectively) before and after ligands treatment. Numbers within the box correspond to the samples’ median *f*_*c*_ values and average diffusion coefficients. The numbers within the box represent the samples’ median *f*_*c*_ values and average diffusion coefficient. All the data are represented as median values with ± SD for *f*_*c*_ (A) and median values with ± SEM for diffusion coefficient (B, C).

Diffusion coefficients for both receptors support the interpretation of the *f*_*c*_ values (Fig. 3A, grey), with reduced mobility observed with EGF and NRG1 treatment compared to in the absence of ligand treatment (Fig. 3B, 3C). EGF ligand treatment slightly reduced the diffusion coefficient of HER4 from 0.34 μm^2^/s to 0.28 μm^2^/s (Fig. 3B) while significantly slowing down EGFR from 0.28 μm^2^/s to 0.17 μm^2^/s (Fig. 3C). EGF may have induced the formation of multimers within EGFR constructs, explaining the higher reduction in receptor mobility with the treatment (Fig. 3C). The NRG1 ligand treatment also significantly reduces the mobility of HER4 proteins (Fig. 3B). NRG1 induces dimerization of HER4 and heteromultimerization between HER4 and EGFR, contributing to the total reduction in protein mobility. The diffusion coefficient of HER4 decreases significantly with NRG1 treatment (Fig. 3B, p < 0.0001) compared to EGF (Fig. 3B, p < 0.01). Like HER4, the mobility of EGFR-mCherry shows the same pattern with EGF treatment compared to NRG1 (Fig. 3C, p < 0.0001).

### HER4-HER2 forms a strong dimer in resting cells and a multimer with ligand stimulation

Our next step was to examine the heterotypic interactions between HER4 and HER2. HER2 has long been proposed to interact with all other members of the HER family in cell signaling(Graus-Porta et al. 1997). To test this heterotypic interaction, we performed PIE-FCCS measurements by co-expressing HER4-eGFP and HER2-mCherry receptors in COS-7 cells before and after ligand stimulation. Single cell PIE-FCCS data and fit parameters can be found in the supplemental information document, and the results are summarized in Fig. 4. Fig. 4A (grey) shows the cross-correlation of HER4 and HER2 in resting cells state with a median *f*_*c*_ value of 0.16. This value supports the interpretation that HER4 interacts directly with HER2 without ligand stimulation. Earlier studies have reported ligand-independent heterodimerization of HER2 and HER3, but to date, there has been no direct experimental investigation of HER4 and HER2 interactions in live cells.

**Figure 4.**
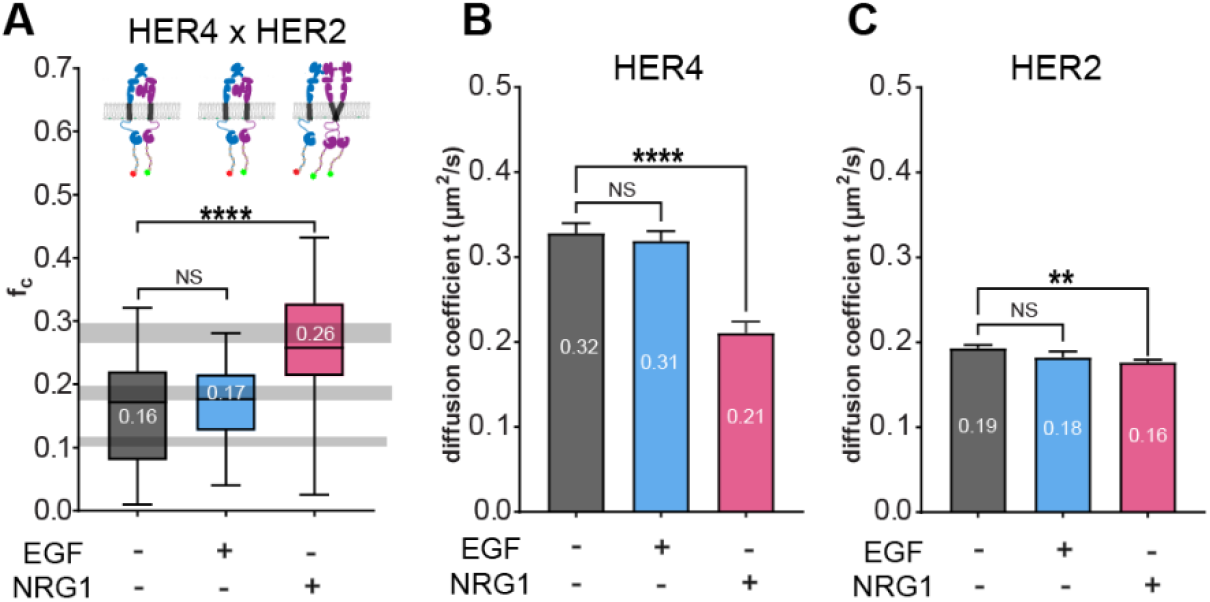
PIE-FCCS measurement of HER4 and HER2 upon ligand stimulation. (A) a fraction of cross-correlation measurement between the HER4 and HER2 proteins co-expressed in COS-7 cells before (-) and after (+) EGF /NRG1 ligand treatment. An illustration of HER proteins cartoon depicted based on PIE-FCCS results in Fig. 4A. (B, C) represent the average diffusion coefficients of protein (HER4 and HER2, respectively) before and after ligands treatment. Numbers within the box correspond to the samples’ median *f*_*c*_ values and average diffusion coefficients. The numbers within the box represent the samples’ median *f*_*c*_ values and average diffusion coefficient. All the data are represented as median values with ± SD for *f*_*c*_ (A) and median values with ± SEM for average diffusion coefficient (B, C).

The median *f*_*c*_ value for the complex remained unchanged following EGF stimulation (*f*_*c*_ = 0.17, Fig. 4A) as well as the diffusion coefficients (Fig. 4B and 4C), which was expected because EGF does not bind HER2 or HER4. However, addition of NRG1 led to an almost two-fold increase of cross-correlation value from 0.16 in resting state to 0.26 after ligand treatment. This increase suggests that NRG1 stimulation drives the HER4-HER2 heterodimer into a larger heteromultimers. This is consistent with recent high-resolution structural data that resolved the ligand-bound HER4-HER2 heteromer(Trenker et al. 2023). HER2 proteins exhibit moderate changes in diffusion coefficient after NRG1 treatment compared to HER4, which may be explained by its lack of homodimerization (p < 0.01, Fig. 4C). One possible explanation for the slow diffusion of HER2 is its extended structure, which could interact with other membrane proteins(Jaulin-Bastard et al. 2001; Jeong et al. 2017). Single cell PIE-FCCS data and fit parameters can be found in Fig. S7 and Table S4).

### NRG1 ligand stimulation drives multimer formation between HER4-HER3 proteins

Next, we measured the degree of interaction between HER4 and HER3 proteins in the absence of ligands and after the addition of EGF and NRG1. The PIE-FCCS experiments were conducted as described above, and single cell PIE-FCCS data and fit parameters can be found in Fig. S8 and Table S5). A median *f*_*c*_ value of 0.12 (Fig. 5A, grey) was observed for HER4 and HER3, indicating ligand-independent heterodimerization. We next stimulated the receptors with EGF or NRG1 ligands using the same concentrations and incubation conditions as above. Following EGF stimulation, we did not detect any change in the *f*_*c*_ value (*f*_*c*_ = 0.11, Fig. 5A, blue), which was expected since EGF is not reported to bind HER3 or HER4. The diffusion coefficients of both receptors after EGF treatment also remained unchanged (Fig. 5B and 5C, blue). We next incubated HER4 and HER3 expressing cells with their specific NRG1 ligand. our PIE-FCCS measurements show a larger median *f*_*c*_ value of 0.22 (Fig. 5A, magenta), indicating their assembly into larger heteromultimers complex. After NRG1 ligand treatment, the average diffusion coefficient of HER4 and HER3 drastically decreased by 42% (0.36 to 0.21 μm^2^/s) and 32% (0.22 to 0.15 μm^2^/s), respectively, which was further evidence of HER3-HER4 hetero-multimerization (Fig. 5B and 5C).

**Figure 5.**
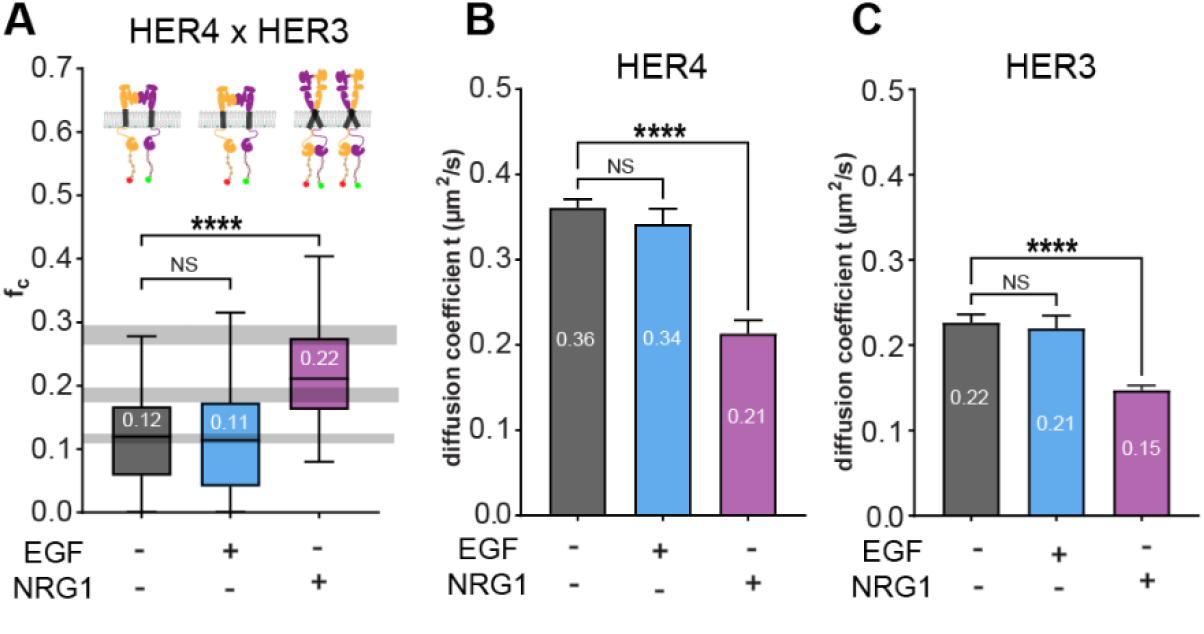
Heterotypic interaction of HER4 and HER3 upon EGF and NRG1 stimulation. (A) our PIE-FCCS measurements reveal the heteromultimerization states for HER4 and HER3 in resting cells. The heteromeric interaction is not affected by EGF ligands, but increases with the HER3/HER4 specific ligand, NRG1. (B, C) The average diffusion coefficients of HER4 and HER3 are shown before and after ligand treatment. The median *f*_*c*_ values and the average diffusion coefficients are displayed inside each box plot.

## Discussion

Communication between cells is as vital in an organism as it is in human relationships. Membrane protein receptors, like a listening ear, receive signals and transmit them inside the cell where the decision-making process develops as a complex network of protein-protein interactions(Lemmon and Schlessinger 2010). Dimerization of RTKs is part of the signal transmission and is therefore integral to cell communication(Lemmon et al. 2014). Understanding how HER proteins are associated with one another is essential for developing more effective therapies and for targeting HER proteins in cancer(Kumagai et al. 2021; Tebbutt et al. 2013). Various organs in the human body exhibit different levels of HER expression, but this does not always account for the large diversity of behaviors. The phosphorylation and dimerization rates of HER depend on the combination of subtypes in pairs, resulting in diverse kinetic rates(Okada et al. 2022).

To better understand the molecular assembly of HER receptors in the cell plasma membrane, we investigated the pair-wise interactions of HER proteins in live cells using PIE-FCCS. This approach enabled us to observe the spatiotemporal assembly of HER family proteins in situ and their response to ligand stimulation. We first investigated the homodimerization state of HER proteins without ligand stimulation (Fig. 1). From this analysis we conclude that EGFR, HER2, and HER4 are predominantly monomeric, while HER3 forms a ligand-independent homodimer. We then investigated heterodimerization between each pair of proteins. We then investigated the pair-wise heterodimerization between each of the four HER-family proteins. Surprisingly, our PIE-FCCS measurements showed that HER4 forms heteromeric interactions with all other HER-family proteins in resting cells. Among the HER4 heterodimers, HER4-HER2, shows the highest degree of cross-correlation, suggesting it is the highest affinity heterodimer in the absence of ligands. EGF ligand stimulation had little to no effect on the HER4 heterodimers, whereas NRG1 induced higher-order hetero-multimers.

The expression level of HER4 is usually low in healthy cells, whereas it is considerably overexpressed in some tumor types and in cancer patients that receive EGFR TKIs(El-Gamal et al. 2021; Segers et al. 2020). HER4 overexpression is often associated with poor prognosis, which supports the hypothesis that HER4-EGFR heterodimers may function as oncoproteins in various cancers(Lucas et al. 2022). An analysis of the HER family gene expression profiles in triple-negative breast cancer (defined by the lack of HER2 expression and estrogen and progesterone receptors), showed that increased HER4 expression was linked to a poor prognosis(Kim et al. 2016). The study suggests that HER4 expression could be used as a marker for predicting response to therapy in triple-negative breast cancer(Kim et al. 2016). Crosstalk between EGFR and HER4 modifies the response to HER4 ligands, indicating that signaling by HER4 homodimers differs from that by HER4-EGFR heterodimers. The heterotypic signaling of HER4-EGFR heterodimers has been observed to be associated with oncogenic phenotypes, such as cell proliferation, migration, invasion, and chemoresistance(Haryuni et al. 2019; Tidcombe et al. 2003). The PIE-FCCS measurements reported here provide evidence for heterodimerization between EGFR and HER4 proteins in resting cells, and a significant increase in heteromultimerization after adding the HER4-specific ligand, NRG1. It is worth noting that the EGFR specific ligand, EGF, had no impact on these heteromeric interactions. Our findings generally agree with previous reports of heterodimerization between HER4 and EGFR.

The prototypical example of RTK heterodimerization is HER2 and HER3, which has been investigated extensively(Diwanji et al. 2021; Graus-Porta et al. 1997; Pryor et al. 2015). HER2 lacks the ability to bind ligands, so its kinase function is only activated when heterodimerized with other members of the HER family(Kiavue et al. 2020). HER3 does not have intrinsic kinase activity(Sierke et al. 1997), and so mainly functions by binding a ligand and then dimerizing with HER2 to activate kinase function(Kol et al. 2014). The role of HER3 homodimerization prior to ligand binding is not fully resolved, however it may serve as a nucleating interaction for binding other HER proteins as observed in our heterodimerization measurements(Berger et al. 2004; Pryor et al. 2015; Steinkamp et al. 2014; Van Lengerich et al. 2017; Váradi et al. 2019). One study found that a HER3-EGFR chimera forms a heteromer with NRG treatment only in the presence of HER2; however, the HER4-EGFR chimera did not require HER2(Berger et al. 2004). Another study suggests that HER3 is an obligatory heteromeric partner due to its inability to homodimerize(Váradi et al. 2019). The size of the HER3 protein heteromer varies based on whether it is stimulated by EGF or neuregulin (NRG). A study using single molecule techniques found that HER3 forms a dimer with EGFR in the presence of NRG, however; it forms higher order oligomers when treated with EGF ligand(Van Lengerich et al. 2017). Another single-molecule study shows that HER2’s heteromeric interaction depends on the ligand(Catapano et al. 2023). In the presence of EGFR-specific ligands (EGF and TGF-a), it forms a heteromer with EGFR; however, HER4-specific ligands induce HER2-HER4 heteromer formation, though the process is slow(Catapano et al. 2023). Our PIE-FCCS measurements of HER2-HER3 heteromeric interaction aligned with several previous reports, where we observed substantial heterodimerization in resting cells (Fig. S10A).

Based on our PIE-FCCS measurements, we conclude that HER4 is a high affinity dimerization partner for all HER-family proteins. To summarize our findings, we propose the following model outlined in Fig.6. Under basal conditions, HER4 is arranged into heterodimers with all other HER family members in resting cells (Fig. 6). These cross-interactions may drive cells to transduce signals and suggests a self-organization principle within these complexes. We have demonstrated that HER signaling can be controlled by ligand binding through the formation of heteromultimers that extend beyond dimers. Our model provides a conceptual framework for future experiments, but additional structural studies are required to elucidate mechanistic details. More work is needed to establish the dimerization interfaces that regulate these interactions and the mechanism by which the heterodimers are activated or inhibited in various liganded states.

**Figure 6.**
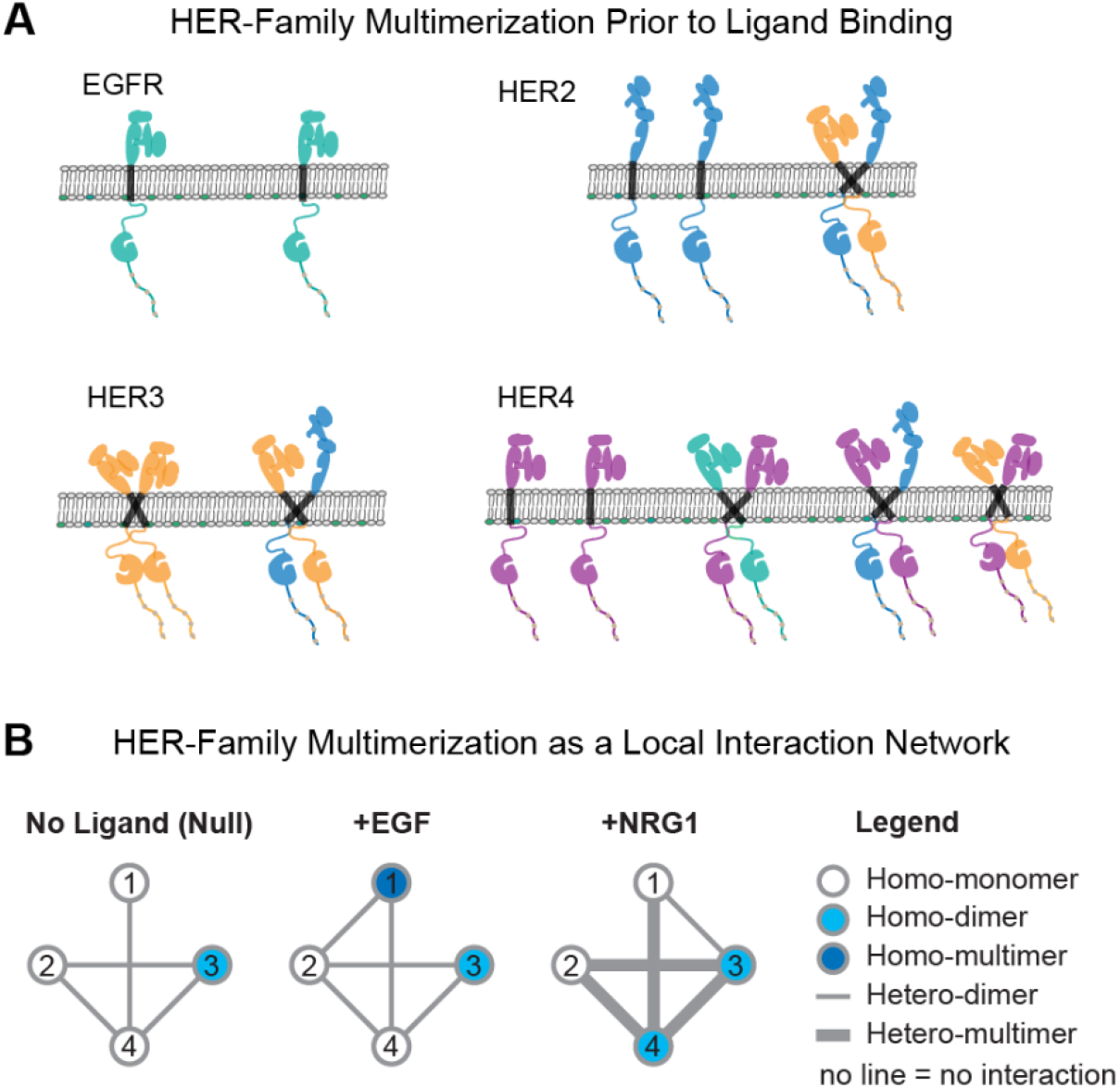
Summary of HER-Family Multimerization. **(A)** Schematic of HER-family multimerization without ligands, where only HER3 (yellow) forms ligand-independent homodimers while EGFR (green), HER2 (blue), and HER4 (magenta) are homo-monomers. HER2 and HER3 assemble into ligand independent heterodimers, while HER4 heterodimerizes with each of the other members: EGFR, HER2, and HER3. **(B)** The multimerization states can be represented as a local network graph, with each node symbolizing a receptor (colored by homo-multimerization state) and each edge (or line) indicating heterodimerization or heteromultimerization. The **No ligand (Null)** graph summarizes the interactions in panel A. When the system is stimulated with EGF, EGFR assembles into homo-multimers and EGFR-HER2 heterodimers. With NRG1 stimulation, HER4 assembles into homodimers, and heterodimerization increases between all of the receptors except EGFR and HER2.

## Materials and Methods

### Cell Culture and Preparation for Imaging

COS-7 cells were used for this study. COS-7 cells were cultured in Dulbecco’s modified Eagle’s medium supplemented with 10% fetal bovine serum and 1% penicillin-streptomycin. Transfection of the plasmids was carried out with 70 to 80% confluent cells in 35 mm glass-bottom MatTek dishes. Plasmids coding EGFR, HER2, HER3 and HER4 were subcloned to eGFP-N1 and mCherry-N1 vectors by XhoI and AgeI digestion. The cells were transiently transfected approximately 20 hours before the data collection with the protein of interest using Lipofectamine 2000 reagent (Thermo Fisher Scientific). A total of 2.5 μg DNA with a 1:1 ratio of mCherry and eGFP-tagged plasmids was used to express both species of fluorescent tagged receptors evenly at a local density of 100–1200 receptors/μm^2^ in the cell measurements reported here (Fig. S9). Before the PIE-FCCS measurements were performed, the media was changed to Opti-MEM I Reduced Serum Medium without phenol red (Thermo Fisher Scientific). For each complex, measurements were taken for both the ligand-free and ligand-stimulated state, using either recombinant human EGF (Sigma-Aldrich, St. Louis, MO) or NRG1 as ligand. In order to stimulate receptor-expressing cells, a stock solution (20 μg/ml) was diluted to 500 ng/ml in Opti-medium (imaging media) and added approximately 15 minutes prior to data collection. Data was collected for a maximum of one hour following stimulation.

### Control Samples

We employed three samples for the alignment of the laser and microscope before data collection. Both 488 and 561 lasers have optical volume differences; therefore, we utilized double-labeled DNA strands to calibrate the volume correction fluorescence value of a sample(Kaliszewski et al. 2018). We applied two samples for the calculation of *f*_*c*_ value of membrane proteins. These negative and positive constructs have a short, lipidated peptide sequence for membrane anchoring and, in the case of GCN4, an α-helical leucine zipper motif for dimerization. From the correlation functions, we obtained the fraction of cross-correlation (*f*_*c*_) and the effective diffusion coefficients of the eGFP and mCH-labeled proteins as described previously. The *f*_*c*_ values are indicative of the co-diffusion of the two receptors (Fig. S2A). The median *f*_*c*_ value of 0.01 for SRC (Fig. S2B, green) and 0.15 for GCN4 (Fig. S2B, blue) are indicative of their monomeric and dimeric state on the plasma membrane. The dimer *f*_*c*_ value of 0.15 is smaller than that observed for a covalent dimer (e.g. duplex DNA, Fig. S2B, red). Our duplex DNA control a 40-nucleotide sequence with <50% G-C content [ACA AGC TGG AGT ACA ACT ACA ACA GCC ACA ACG TCT ATA T] was labeled with carboxy-tetramethylrhodamine (TAMRA) at the 5’ end and 6-carboxyfluorescein (FAM) at the 3’ end (Integrated DNA Technologies). An excess of the unlabeled strand was used to anneal the synthesized complementary strand pair as per the supplier’s protocols. The double-stranded DNA, labeled with both TAMRA and FAM (TAMRA-40-FAM), was diluted to a final concentration of 100 nM using 10 mM TE buffer. For the 3D sample data collection, laser powers were set to 7 μW and 7 μW for the 488 nm and 561 nm lasers, respectively.

### Data Collection and Analysis

The time-time autocorrelation function is employed for single color FCS to analyze intensity fluctuations. The results were plotted on a semi-log axis to better view the time scale. In principle, ACF amplitude is inversely proportional to the average number of molecules in the observation area. In FCCS, two fluorescent probes are used to analyze the emission independently of one another using two separate spectrally distinct probes. As a result, both populations have a corresponding ACF, and molecular density can be determined independently. With PIE-FCCS, two laser pulses are phase-delayed calculating the exact arrival time of each laser pulse’s emission photons.

The FCCS data were recorded using pulsed interleaved excitation and time-correlated single-photon detection with a custom inverted microscope setup. A supercontinuum pulsed laser (9.2 MHz repetition rate, SuperK NKT Photonics, Birkerød, Denmark) was split into two beams of 488 and 561 nm using a series of filter and mirror combinations. In order to achieve PIE, the beams are directed through separate optical fibers of varying lengths, causing a delay in arrival time between them. This eliminates spectral crosstalk between the detectors. Before entering the microscope, the beams were overlapped by a dichroic beam splitter (LM01-503-25, Semrock) and a customized filter block (zt488/561rpc, zet488/561m, Chroma Technology). The overlapping beams of light were focused by the objective (×100 TIRF) to a limited diffraction spot on the peripheral membrane of a Cos-7 cell expressing the receptor constructs. In time-tagged time-resolved mode, photons were detected by individual avalanche photodiodes (Micro-Photon Devices). In order to verify the alignment of the system, including the overlap of a confocal volume, a short fluorescently tagged DNA fragment was used. Prior to the experimental samples, positive and negative controls (Fig. S2, S3) were tested to compare the fit parameters.

The excitation beams were focused on the peripheral membranes to measure only membrane-bound receptors. Only the cell’s flat, peripheral membrane area was scanned to prevent fluorescence from cytosolic organelles or vesicles. The data collection was performed on one area of the membrane in each sample. Each area was assessed six times, with each acquisition lasting 10 seconds. MATLAB scripts were used to calculate each sample’s auto- and cross-correlation curves. As described in the previous work, we fit a single component, the 2D diffusion model, to the averaged curves of six consecutive acquisitions per area.

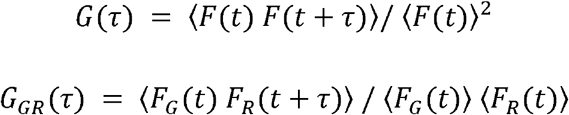

The intensity fluctuations were gated using PIE prior to cross-correlation and autocorrelation analyses. The intensity at time F(t) was compared to the intensity at a later time F(t + ⍰) in an autocorrelation analysis. As a function of time, self-similarity allowed for the interpretation of quantitative information, such as diffusion and particle number. In Equation 1, the intensity fluctuations were divided into 10-second bins and normalized to the square of the average intensity. The autocorrelation functions were fit to the following model for two-dimensional diffusion in the membrane that accounts for triplet relaxation and dark state dynamics.

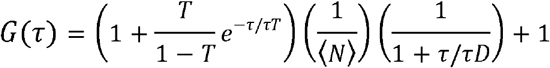

The diffusion coefficient of the sample has been calculated by utilizing the standard formula given below.

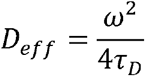

Finally, the cross-correlation function (fraction of correlation) was by using the autocorrelation function of both fluorophores.

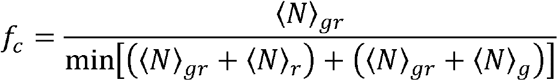

## Acknowledgements

This work was supported by the National Science Foundation grant CHE-1753060, the Human Frontiers of Science Program No. RGP0059/2019, and the American Lung Association Lung Cancer Discovery Award, LCD-1035035.

## Author contributions

Conceptualization: AWS, SK, PKS; Plasmid construction: SK, PKS; Performed Experiments: PKS, SK; Data Analysis: PKS, SK, AWS, Writing original draft: PKS, AWS, Writing-Review and editing: AWS, PKS, SK Supervision and Project Administration: AWS, Funding Acquisition: AWS

