## Supplemental Material for "HER4 is a high affinity dimerization partner for all EGFR/HER/ErbB-family proteins"

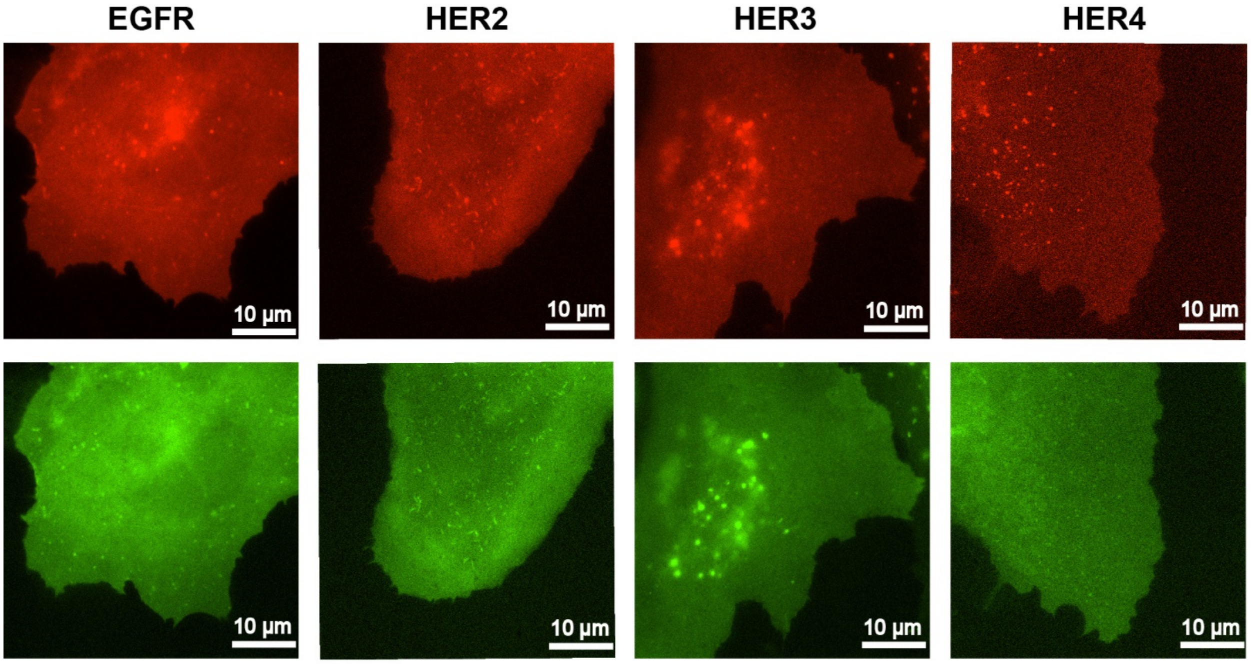


**Supplemental Figure S1.**

Epifluorescence images of COS-7 cells expressing HER family proteins (mCherry and eGFP tagged) were used to study homodimer and heteromeric interactions. Unless otherwise noted, these HER protein expression on the plasma membrane of COS-7 cells is recorded in the resting cell conditions.

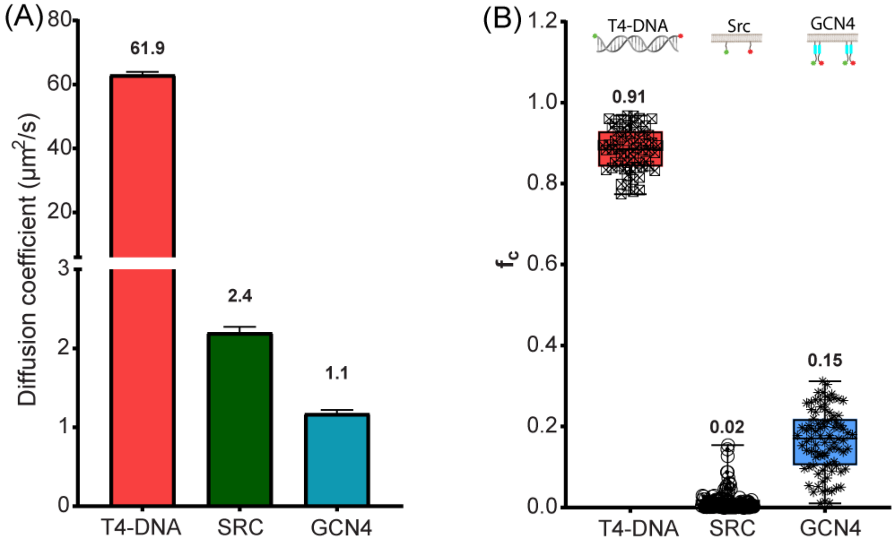


**Supplemental Figure S2.**

The diffusion coefficients and cross-correlation of control samples were employed for the calibration of microscope alignment. Panel A shows the diffusion coefficient of all three control samples extracted after data analysis and fitting. Panel B represents the fraction of correlation ($f_c$) of monomeric (SRC) and dimeric (GCN4) protein construct expressed on the plasma membrane in COS-7 cells, respectively, and double-labeled (two fluorophores) 3D sample of DNA. Panel B also shows the sketch of all three control samples.

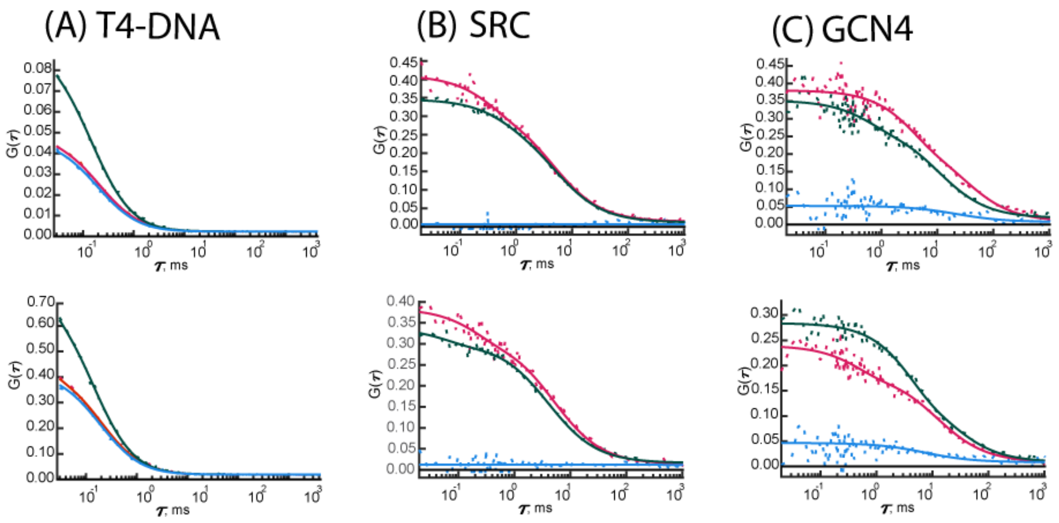


**Supplemental Figure S3.**

Representative PIE-FCCS graphs of control samples used for microscope alignment before data collection. Fig. S3 A shows the diffusion profile of a 3D sample of the double-labeled DNA used to calibrate the microscope. Fig. B and C represent the diffusion profile of monomeric (SRC) and dimeric (GCN4) protein construct expressed on the plasma membrane in COS-7 cells, respectively. In each graph, the green line is the autocorrelation function (ACF) for green fluorophore, the red line is the ACF for red fluorophore, and the blue line is the cross-correlation function (CCF) between both green and red.

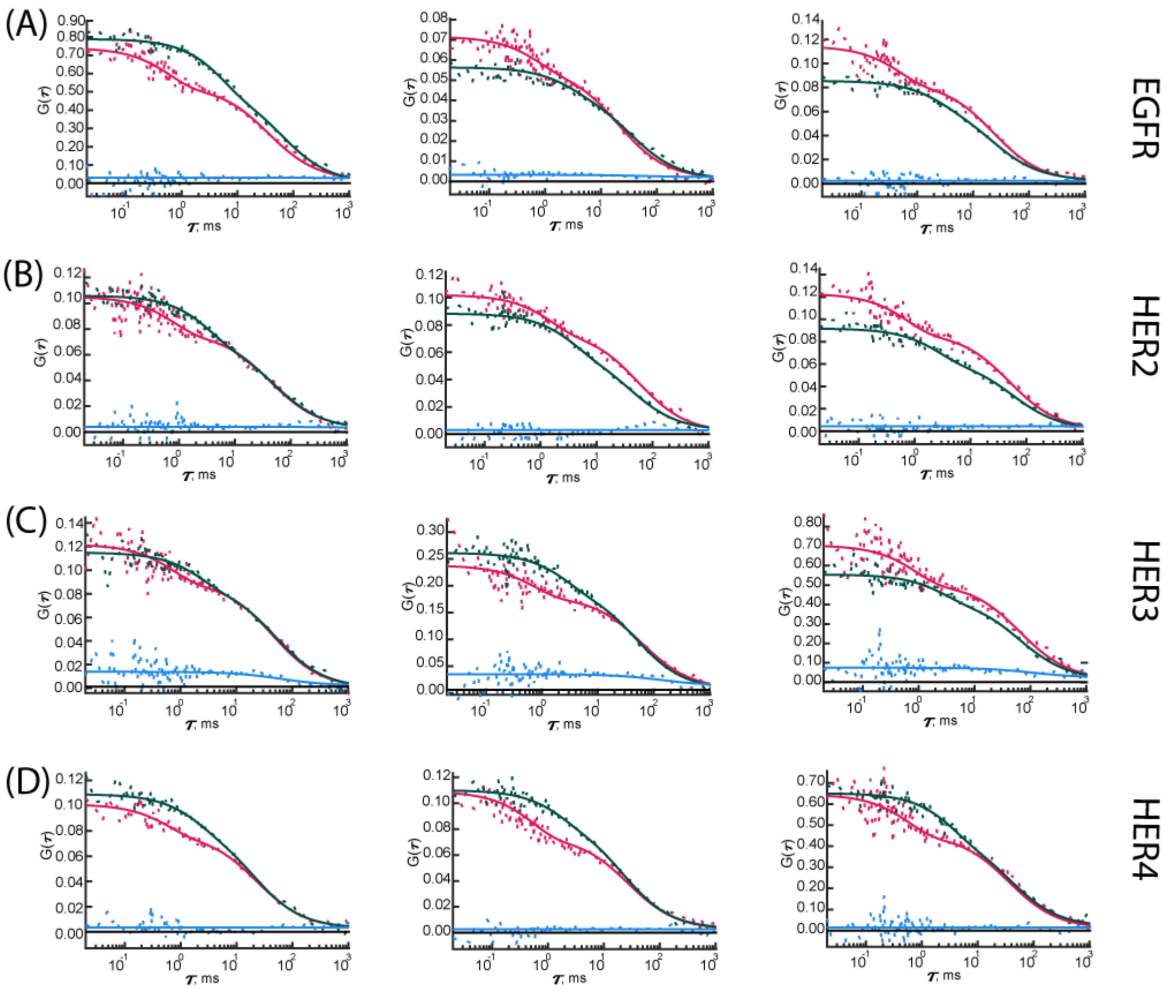


**Supplemental Figure S4.**

PIE-FCCS graphs homodimers of all four HER family members on the plasma membrane of COS-7 cells. Fig S3 A, B, C, and D show the diffusion profile of EGFR, HER2, HER3, and HER4 homo-oligomers in resting or untreated cell conditions, respectively. In each graph, the green line is the autocorrelation function (ACF) for green fluorophore, the red line is the ACF for red fluorophore, and the blue line is the cross-correlation function (CCF) between both green and red.

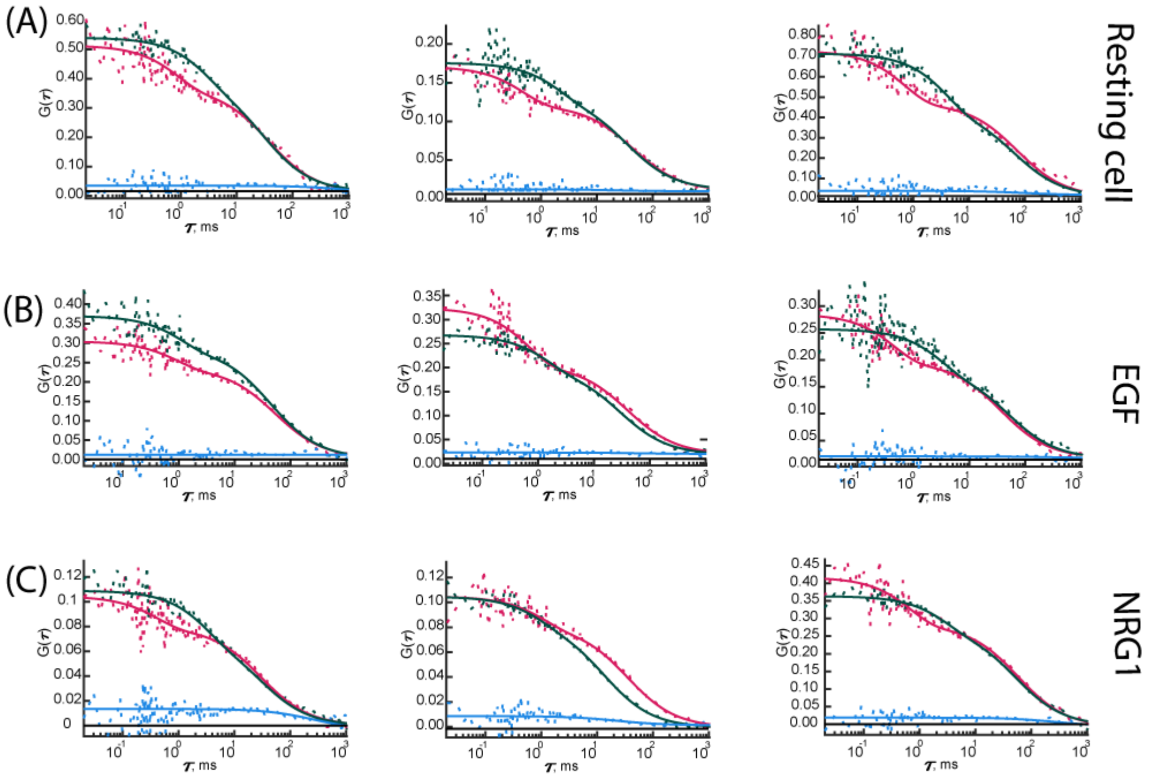


**Supplemental Figure S5.**

PIE-FCCS graphs of diffusion profile of HER4 homodimers on the plasma membrane of COS-7 cells. Fig S5 A shows the diffusion profile of HER4 in resting or untreated cell conditions. Fig S5 B and C show the diffusion profile of HER4 with ligands treatment, EGF and NRG1 respectively. In each graph, the green line is the autocorrelation function (ACF) for green fluorophore, the red line is the ACF for red fluorophore, and the blue line is the cross-correlation function (CCF) between both green and red.

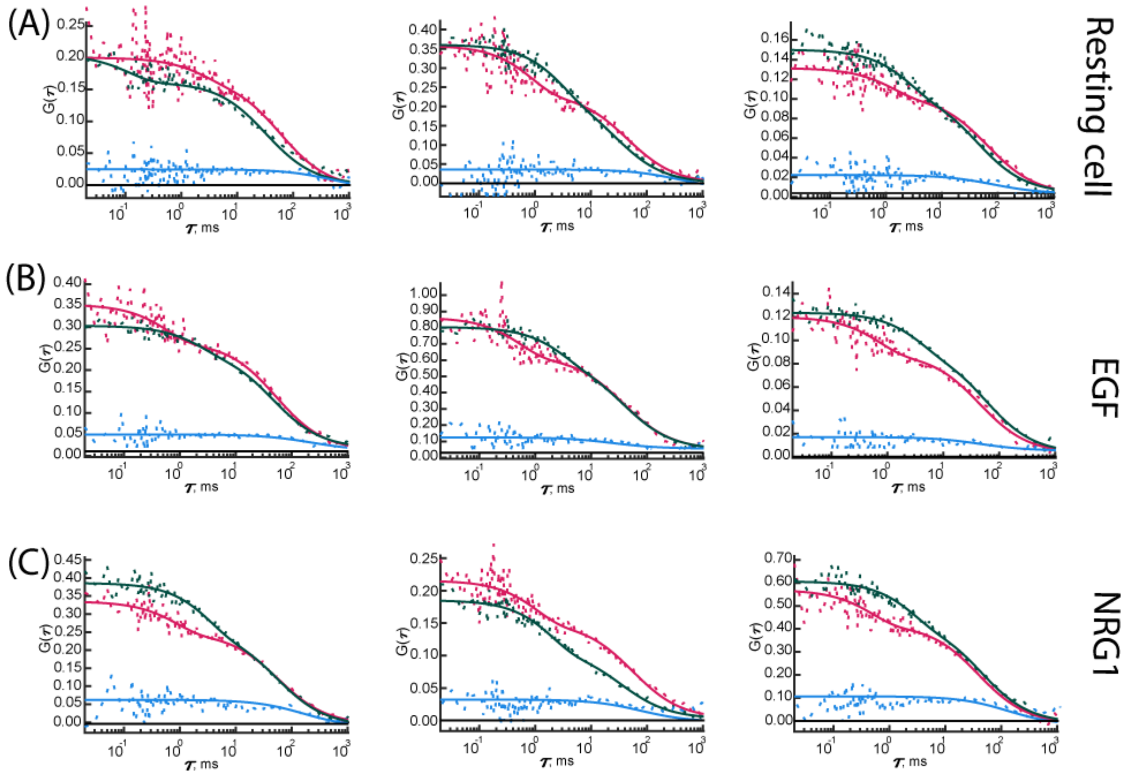


**Supplemental Figure S6.**

PIE-FCCS graphs of diffusion profile of EGFR and HER4 heterodimers on the plasma membrane of COS-7 cells. Fig S6 A shows the diffusion profile of EGFR and HER4 proteins in resting or untreated cell conditions. Fig S6 B and C show the diffusion profile of EGFR and HER4 proteins with ligands treatment, EGF and NRG1 respectively. In each graph, the green line is the autocorrelation function (ACF) for green fluorophore, the red line is the ACF for red fluorophore, and the blue line is the cross-correlation function (CCF) between both green and red.

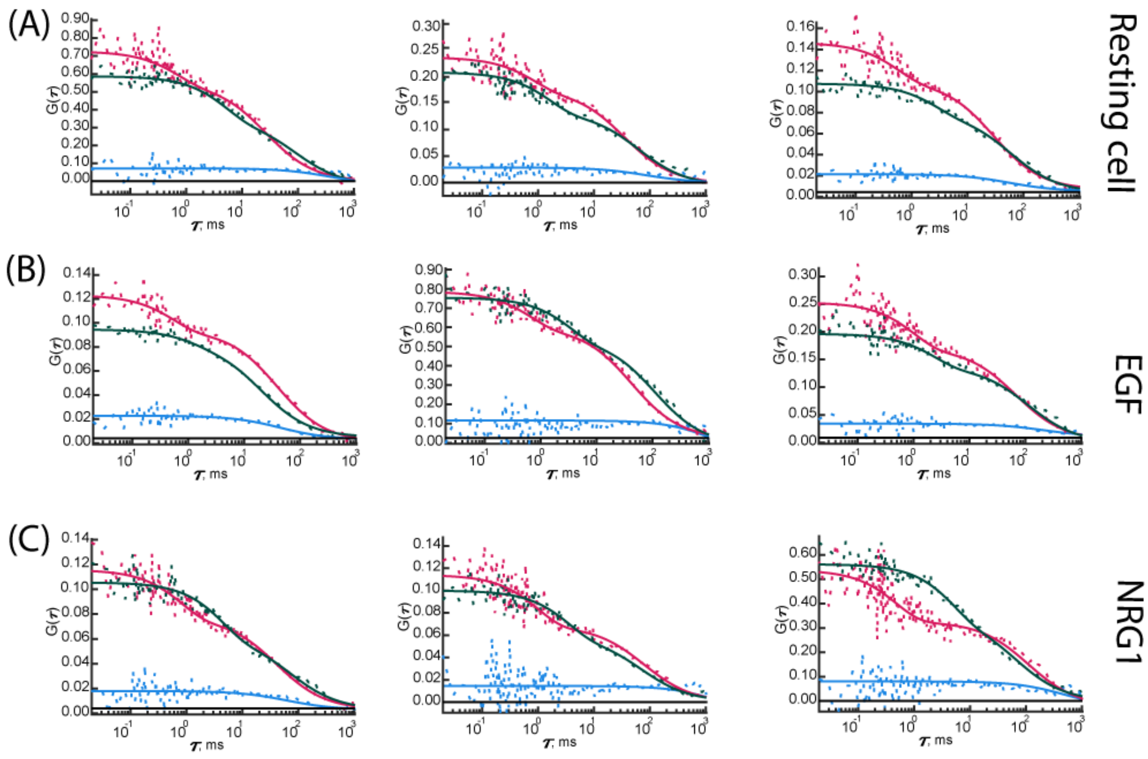


**Supplemental Figure S7.**

PIE-FCCS graphs of diffusion profile of HER2 and HER4 heterodimers on the plasma membrane of COS-7 cells. Fig S7 A shows the diffusion profile of HER2 and HER4 proteins in resting or untreated cell conditions. Fig S7 B and C show the diffusion profile of HER2 and HER4 proteins with ligands treatment, EGF and NRG1 respectively. In each graph, the green line is the autocorrelation function (ACF) for green fluorophore, the red line is the ACF for red fluorophore, and the blue line is the cross-correlation function (CCF) between both green and red.

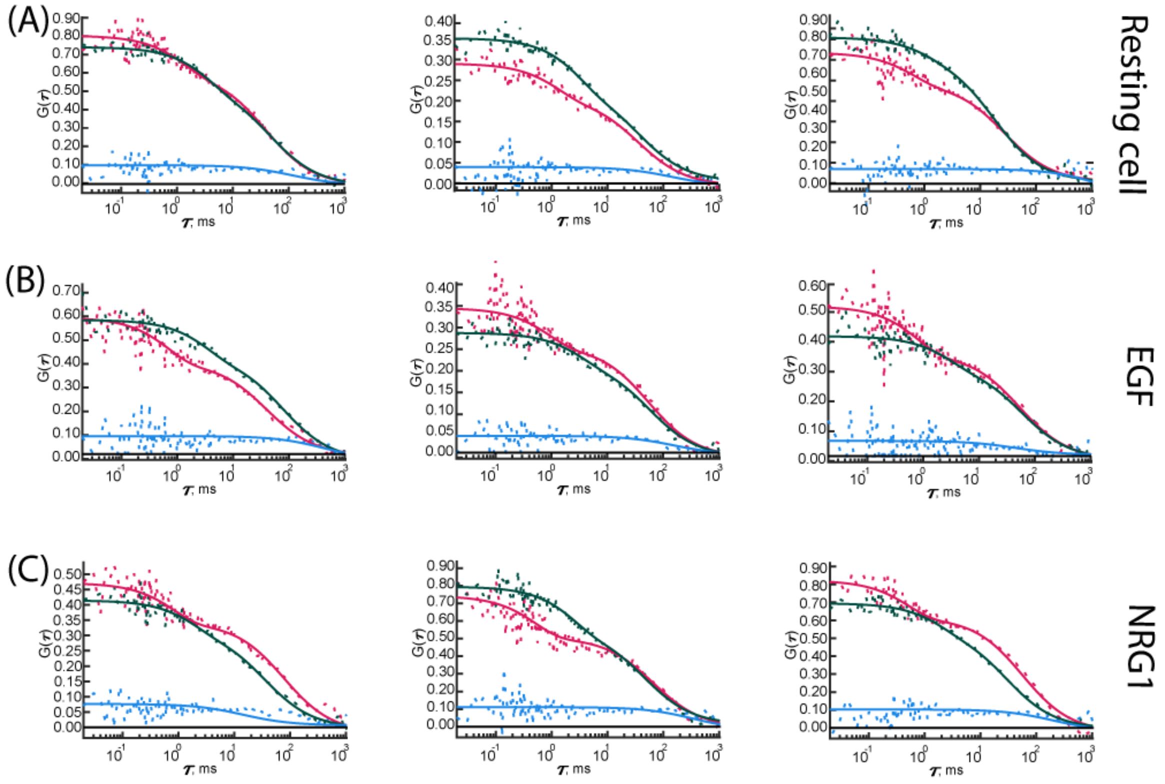


**Supplemental Figure S8.**

PIE-FCCS graphs of diffusion profile of HER3 and HER4 heterodimers on the plasma membrane of COS-7 cells. Fig S8 A shows the diffusion profile of HER3 and HER4 proteins in resting or untreated cell conditions. Fig S8 B and C show the diffusion profile of HER2 and HER4 proteins with ligands treatment, EGF and NRG1 respectively. In each graph, the green line is the autocorrelation function (ACF) for green fluorophore, the red line is the ACF for red fluorophore, and the blue line is the cross-correlation function (CCF) between both green and red.

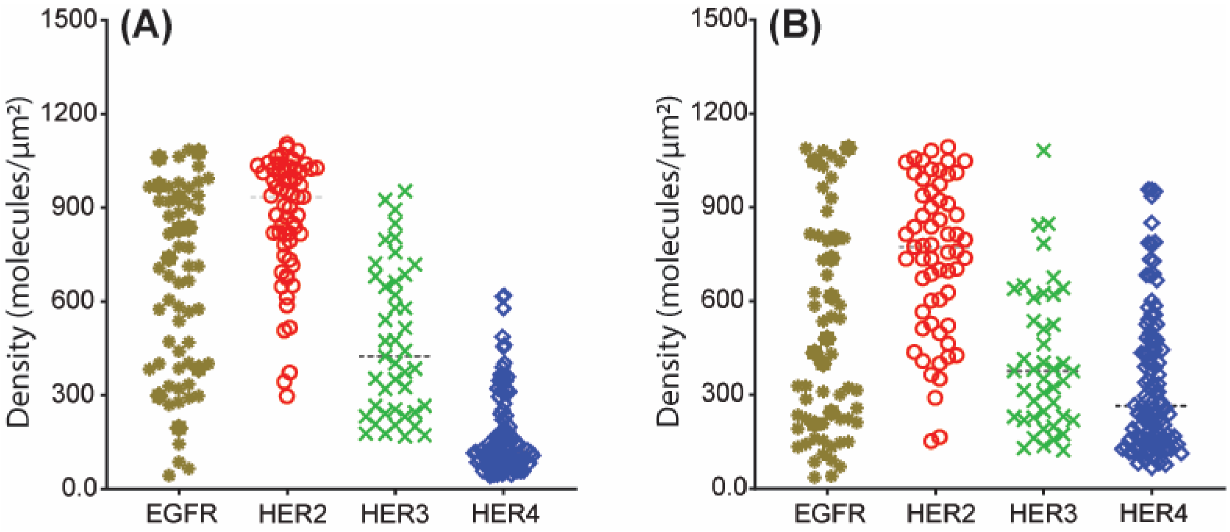


**Supplemental Figure S9.**
The molecular densities of HER family proteins COS-7 cells expressing during these heteromeric interactions studies data collection. Panels A and B show the molecules density of mCherry and eGFP tagged protein, respectively.

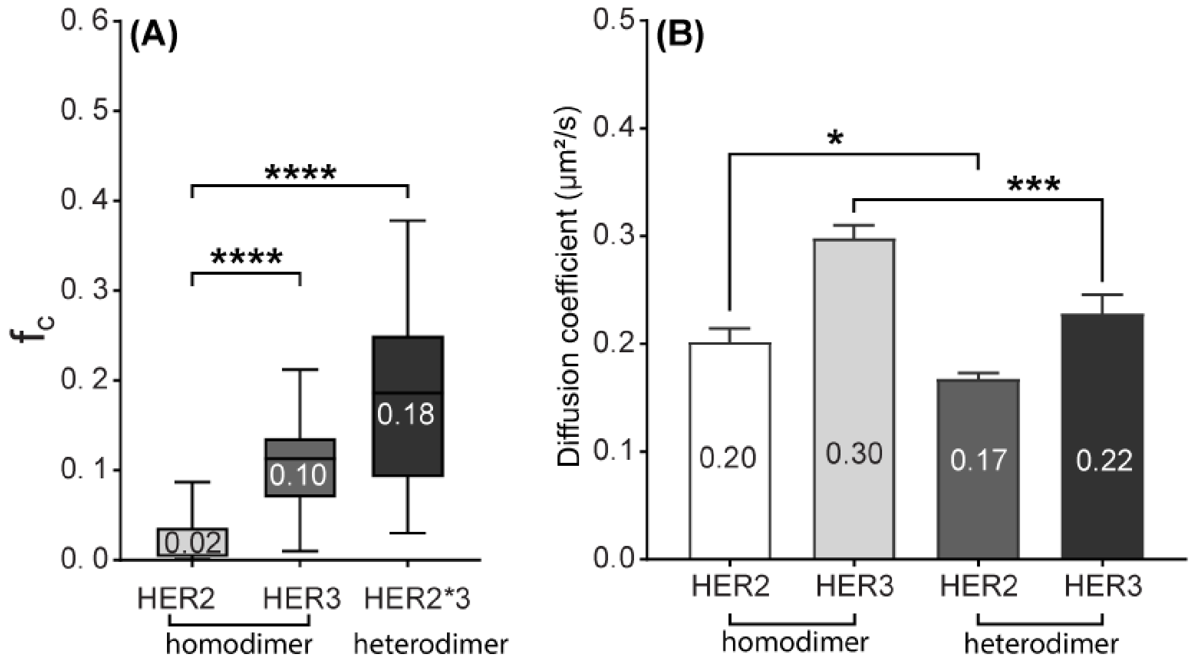


**Supplemental Figure S10.**

The fraction of correlation and diffusion coefficient of HER 2 and HER3 proteins on the plasma membrane of COS-7 cells. (A) The average median $f_c$ value of HER2 (light gray), HER3 (medium gray), and cross-interaction of both proteins in resting cells (homotypic). In resting or non-stimulating conditions, HER2 protein stayed monomer while the HER3 proteins were present in their dimeric form. (B) The receptors' molecular diffusion coefficients also support the average median $f_c$ value and show a reduction in protein mobility during HER2 and HER3 heterodimer formation as compared to monomeric states.

**Table S1.**

Summary of experimental parameters obtained from PIE-FCCS measurements for HER proteins (Homodimerization) in COS-7 cells membranes. Data values represent mean ± SD (n > 60) and are rounded to the nearest integer. While the fraction correlated ($f_c$) values are reported as mean, and median measurements, only the median values are plotted in figures.

| Parameter | HER receptors homodimerization in COS-7 cell membranes | | | |
|---|---|---|---|---|
| | **EGFR** | **HER2** | **HER3** | **HER4** |
| Data points | 60 | 61 | 62 | 111 |
| $f_c$ (median) | 0.02 | 0.01 | 0.10 | 0.02 |
| $f_c$ (mean) | 0.02±0.02 | 0.02±0.02 | 0.10±0.05 | 0.02±0.05 |
| Density (R)[a] (mol/$\mu m^2$) | 641±299 | 865±193 | 472±239 | 170±128 |
| Density (G)[b] (mol/$\mu m^2$) | 483±336 | 741±243 | 410±226 | 338±231 |
| $D_R$ ($\mu m^2 \cdot s^{-1}$)[c] | 0.38±0.07 | 0.23±0.10 | 0.29±0.08 | 0.37±0.07 |
| $D_G$ ($\mu m^2 \cdot s^{-1}$)[d] | 0.31±0.20 | 0.18±0.06 | 0.23±0.10 | 0.38±0.07 |
| Brightness (R) | 361±70 | 304±57 | 332±59 | 356±89 |
| Brightness (G) | 562±136 | 448±90 | 532±85 | 542±199 |

***a,*** Density of receptors in the red channel. ***b,*** Density of receptors in the green channel. ***c***, Diffusion of receptors in the red channel. ***d***, Diffusion of receptors in the green channel.

**Table S2.**

Summary of experimental parameters obtained from PIE-FCCS measurements for HER4 protein homodimerization in COS-7 cells membranes before and after ligands treatments. Data values represent mean ± SD (n > 60) and are rounded to the nearest integer. While the fraction correlated ($f_c$) values are reported as mean, and median measurements, only the median values are plotted in figures.

| **Parameter** | **HER4 receptors homodimerization with ligands treatment in COS-7 cell membranes** | | |
|---|---|---|---|
| | **No ligands** | **EGF** | **NRG1** |
| Data points | 111 | 72 | 62 |
| $f_c$ (median) | 0.02 | 0.02 | 0.12 |
| $f_c$ (mean) | 0.02±0.05 | 0.03±0.04 | 0.12±0.06 |
| Density (R)$^a$ (mol/μm$^2$) | 170±128 | 251±189 | 151±82 |
| Density (G)$^b$ (mol/μm$^2$) | 338±231 | 139±113 | 227±189 |
| $D_R$ (μm$^2$ · s$^{-1}$)$^c$ | 0.37±0.07 | 0.36±0.06 | 0.24±0.05 |
| $D_G$ (μm$^2$ · s$^{-1}$)$^d$ | 0.38±0.07 | 0.37±0.07 | 0.25±0.10 |
| Brightness (R) | 356±89 | 390±86 | 340±64 |
| Brightness (G) | 542±200 | 515±199 | 551±149 |

***a,*** Density of receptors in the red channel. ***b,*** Density of receptors in the green channel. ***c***, Diffusion of receptors in the red channel. ***d***, Diffusion of receptors in the green channel.

**Table S3.**

Summary of experimental parameters obtained from PIE-FCCS measurements for EGFR and HER4 protein heterodimerization in COS-7 cells membranes before and after ligands treatments. Data values represent mean ± SD (n > 60) and are rounded to the nearest integer. While the fraction correlated ($f_c$) values are reported as mean, and median measurements, only the median values are plotted in figures.

| **Parameter** | **EGFR and HER4 protein heterodimerization before and after ligands treatment in COS-7 cell membranes** | | |
|---|---|---|---|
| | **No ligands** | **EGF** | **NRG1** |
| Data points | 81 | 62 | 60 |
| $f_c$ (median) | 0.12 | 0.17 | 0.21 |
| $f_c$ (mean) | 0.12±0.05 | 0.18±0.05 | 0.21±0.08 |
| Density (R)$^{a}$ (mol/μm$^2$) | 219±216 | 224±128 | 361±216 |
| Density (G)$^{b}$ (mol/μm$^2$) | 268±218 | 244±148 | 194±115 |
| $D_R$ (μm$^2$ · s$^{-1}$)$^{c}$ | 0.35±0.07 | 0.29±0.07 | 0.24±0.11 |
| $D_G$ (μm$^2$ · s$^{-1}$)$^{d}$ | 0.33±0.19 | 0.17±0.05 | 0.25±0.17 |
| Brightness (R) | 342±93 | 334±77 | 459±137 |
| Brightness (G) | 567±164 | 636±184 | 450±136 |

***a,*** Density of receptors in the red channel. ***b,*** Density of receptors in the green channel. ***c***, Diffusion of receptors in the red channel. ***d***, Diffusion of receptors in the green channel.

**Table S4.**

Summary of experimental parameters obtained from PIE-FCCS measurements for HER2 and HER4 protein heterodimerization in COS-7 cells membranes before and after ligands treatments. Data values represent mean ± SD (n > 60) and are rounded to the nearest integer. While the fraction correlated ($f_c$) values are reported as mean, and median measurements, only the median values are plotted in figures.

| **Parameter** | **HER2 and HER4 protein heterodimerization before and after ligands treatment in COS-7 cell membranes** | | |
|---|---|---|---|
| | **No ligands** | **EGF** | **NRG1** |
| Data points | 94 | 76 | 62 |
| $f_c$ (median) | 0.17 | 0.18 | 0.26 |
| $f_c$ (mean) | 0.16±0.08 | 0.17±0.06 | 0.26±0.09 |
| Density (R)$^a$ (mol/μm$^2$) | 901±537 | 973±551 | 678±464 |
| Density (G)$^b$ (mol/μm$^2$) | 183±157 | 224±181 | 205±147 |
| $D_R$ (μm$^2$ · s$^{-1}$)$^c$ | 0.33±0.11 | 0.32±0.07 | 0.17±0.03 |
| $D_G$ (μm$^2$ · s$^{-1}$)$^d$ | 0.24±0.12 | 0.18±0.05 | 0.21±0.11 |
| Brightness (R) | 329±96 | 347±87 | 356±126 |
| Brightness (G) | 530±174 | 500±102 | 491±187 |

***a,*** Density of receptors in the red channel. ***b,*** Density of receptors in the green channel. ***c***, Diffusion of receptors in the red channel. ***d***, Diffusion of receptors in the green channel.

**Table S5.**

Summary of experimental parameters obtained from PIE-FCCS measurements for HER3 and HER4 protein heterodimerization in COS-7 cells membranes before and after ligands treatments. Data values represent mean ± SD (n > 60) and are rounded to the nearest integer. While the fraction correlated ($f_c$) values are reported as mean, and median measurements, only the median values are plotted in figures.

| **Parameter** | **HER3 and HER4 protein heterodimerization before and after ligands treatment in COS-7 cell membranes** | | |
|---|---|---|---|
| | **No ligands** | **EGF** | **NRG1** |
| Data points | 104 | 75 | 65 |
| $f_c$ (median) | 0.12 | 0.11 | 0.21 |
| $f_c$ (mean) | 0.11±0.07 | 0.11±0.08 | 0.22±0.07 |
| Density (R)[a] (mol/$\mu m^2$) | 358±224 | 376±198 | 228±142 |
| Density (G)[b] (mol/$\mu m^2$) | 196±135 | 150±117 | 219±108 |
| $D_R$ ($\mu m^2 \cdot s^{-1}$)[c] | 0.34±0.10 | 0.34±0.12 | 0.15±0.05 |
| $D_G$ ($\mu m^2 \cdot s^{-1}$)[d] | 0.27±0.14 | 0.22±0.11 | 0.21±0.12 |
| Brightness (R) | 363±95 | 334±76 | 351±94 |
| Brightness (G) | 622±151 | 608±151 | 518±184 |

***a,*** Density of receptors in the red channel. ***b,*** Density of receptors in the green channel. ***c***, Diffusion of receptors in the red channel. ***d***, Diffusion of receptors in the green channel.
